# Involvement of the *Pseudomonas aeruginosa* MexAB-OprM efflux pump in the secretion of the metallophore pseudopaline

**DOI:** 10.1101/2020.05.13.092411

**Authors:** Nicolas Oswaldo Gomez, Alexandre Tetard, Laurent Ouerdane, Clémentine Laffont, Catherine Brutesco, Geneviève Ball, Ryszard Lobinski, Yann Denis, Patrick Plésiat, Catherine Llanes, Pascal Arnoux, Romé Voulhoux

## Abstract

The ability for all organisms to acquire metals from their environment is essential for life. To overcome the metal restriction imposed by the host’s nutritional immunity, bacterial pathogens exploits the use of small high metal affinity molecules called metallophores. Metallophores are first synthetized in the cytoplasm, then secreted into the medium where they sequester the metal. The metal-metallophore complex is then imported into the bacterium following binding to dedicated cell surface receptors. Recently, a new family of metallophores has been discovered in pathogenic bacteria called staphylopine in *Staphylococcus aureus* and pseudopaline in *Pseudomonas aeruginosa*. Here, we are expending the molecular understanding of pseudopaline secretion/recovery cycle across the double-membraned envelope of the Gram-negative bacterium *Pseudomonas aeruginosa*. We first revealed that pseudopaline is secreted in a two-step process including export across the inner membrane by the CntI exporter followed by a specific transport across the outer membrane by the MexAB-OprM efflux pump. Such involvement of MexAB-OprM in pseudopaline secretion, reveal a new natural function that extends its spectrum of functions and therefore reasserts its interest as antibacterial target. We then addressed the fate of the recovered metal-loaded pseudopaline by combining *in vitro* reconstitution experiments using radio-labeled pseudopaline subjected to bacterial lysates, and *in vivo* phenotyping in absence of pseudopaline transporters. Our data support the existence of a pseudopaline degradation/modification mechanism, possibly involved in metal release following pseudopaline recovery. All together our data allowed us to provide an improved molecular model of secretion, recovery and fate of this important metallophore by *P. aeruginosa*.

**IMPORTANCE:** Pseudopaline is a broad spectrum metallophore produced and used by *Pseudomonas aeruginosa* to supply the bacterium in metal in metal scarce environments. Here we are investigating the pseudopaline transport/recovery cycle across the bacterial envelope. We are first demonstrating that pseudopaline secretion in the medium is achieved by a specific efflux pump, usually dedicated to the release of toxic compounds such as antibiotics, thus revealing a new natural function for this efflux pump reasserting its interest as antibacterial target. Additional experiments also revealing the existence of an intracellular pseudopaline degradation mechanism providing new clues to another obscure step of the pseudopaline cycle which is the intracellular metal liberation from the imported metal-pseudopaline complex. All together our data allowed us to disclose important aspects of the secretion, recovery and fate of this essential molecule used by *P. aeruginosa* to survive during infections thus constituting new potential targets for antibacterial development.

## INTRODUCTION

The availability of a diversity of metals is essential to support a variety of essential biological functions and bacteria have evolved several mechanisms for their acquisition from environmental sources. For this purpose, bacteria deploy multiple systems that govern both influx and efflux of transition metals allowing cells to maintain a meticulously regulated pool of metals in any circumstances. This is particularly relevant for pathogenic bacteria within a host, where the concentration of free metals is kept extremely low by the host’s nutritional immunity framework, which has very likely evolved to limit the growth of microorganisms in infected tissues (1, 2). Under such metal scarce conditions, the most common strategy used by pathogenic bacteria to import metals involves the synthesis and release of metallophores followed by the capture and import of the metal-metallophore complex (3).

*Pseudomonas aeruginosa* is a versatile opportunistic human pathogen, well equipped for thriving in particularly metal limited environments (4, 5). Indeed, this pathogen is able to survive under strong metal scarcity due to its ability to produce various metallophores. To date, our knowledge of the metallophore-mediated metal uptake capabilities of *P. aeruginosa* has been limited to two iron-specific siderophores, pyochelin and pyoverdine (6) and a recently discovered broad-spectrum metallophore named pseudopaline (7). Pseudopaline is distantly related to plant nicotianamine and belongs to the same opine-type broad-spectrum metallophore family as staphylopine and yersinopine synthetized by *Staphylococcus aureus* and *Yersinia pestis*, respectively (8-10). The *cntOLMI* genes of *P. aeruginosa* encode for the proteins involved in pseudopaline biosynthesis and transport (7). They all cluster together in a single operon negatively regulated by zinc through the Zur repressor (7, 11). The first gene of the operon, *cntO* encodes for the OM component CntO, belonging to the TonB-Dependent Transporter (TBDT) family dedicated to pseudopaline recovery from the environment (7, 12, 13). The second and third genes of the operon, respectively *cntL* and *cntM*, encode the two cytoplasmic soluble enzymes CntL and CntM responsible for pseudopaline biosynthesis in a two steps process. The first step, in which CntL produces the reaction intermediate yNA using S-adenosine methionine (SAM) and L-histidine as substrates, is followed by a NADH reductive condensation of the yNA intermediate with a molecule of α-ketoglutarate (αKG), catalyzed by CntM to produce pseudopaline (7). The fourth gene of the *cnt* operon, *cntI*, encodes the IM protein CntI belonging to the EamA or drug/metabolite transporter (DMT) family involved in pseudopaline export from the cytoplasm to the periplasm after biosynthesis (7).

CntI and CntO were shown to be essential for *P. aeruginosa* growth in Airway Mucus Secretion (AMS) (4) and *P. aeruginosa*’s capacity to colonize mice (12) respectively. Moreover, several transcriptomic analyses showed that *cnt* genes are up-regulated during lung infection (14, 15). During chronic infection, polymorphisms within the promoter sequence of the *cnt* operon were reported, leading to an altered Zur binding site and a less stringent repression (16). Altogether, these pieces of information suggest that the involvement of the Cnt machinery in zinc uptake in a scarce and chelated environment is crucial for *P. aeruginosa* survival during lung infections of cystic fibrosis patients.

Nonetheless, the complete pseudopaline export and recovery pathway remains to be fully elucidated. In particular it remains unknown how pseudopaline crosses the OM to be released in the extracellular milieu. Several other metallophores rely on efflux pumps for their secretion through the OM including the pyoverdine with PvdRT-OpmQ in *P. aeruginosa* (17, 18) or enterobactin with the modular AcrAB-, AcrAD-, and MdtBC-TolC RND efflux pump in *Escherichia coli* (19).

In this study we are completing the understanding of the pseudopaline secretion pathway by demonstrating that, following its export across the IM by CntI, its translocation across the OM is accomplished by the tripartite RND efflux pumps MexAB-OprM, thus revealing the existence of a two-step pseudopaline secretion system in *P. aeruginosa*. We have also observed that the impaired secretion of pseudopaline in absence of its dedicated efflux pump does not lead to periplasmic accumulation of pseudopaline in contrast to the cytoplasmic accumulation observed in absence of IM exporter. We are providing experimental evidence supporting the existence of a pseudopaline modification mechanism in the periplasm. This unidentified pseudopaline modification mechanism may also be involved in physiological pseudopaline alteration, leading to the release of the cognate metal following its recovery from the extracellular medium by CntO.

## RESULTS

### Pseudopaline is required for normal growth, biofilm formation and full extracellular protease activity in minimal chelated media

Pseudopaline is essential for *P. aeruginosa* growth in Airway Mucus Secretion (AMS) and provides an advantage for mice colonization (4, 12). Several transcriptomic approaches showed that the pseudopaline biosynthesis and transport machinery, encoded by the *cnt* operon, is upregulated by *P. aeruginosa* in the lungs of Cystic Fibrosis (CF) patients (14, 15). Similarly, growth in a minimal chelated medium (minimal succinate medium supplemented with 100µM EDTA, MCM) induces a significant expression of the *cnt* operon compared to the LB nutrient rich medium (figure S1). We previously showed that pseudopaline was required for zinc uptake in MCM (7). Altogether, these pieces of information indicate that during infection and growth in MCM, *P. aeruginosa* is starved for Zn, further validating MCM as a medium mimicking the host Zn shortage condition. In order to evaluate the implication of pseudopaline in physiological functions relevant for *P. aeruginosa*’s pathogenesis, we compared different fitness parameters between a wild type and a Δ*cntL* strain, deficient for pseudopaline biosynthesis, during growth in MCM. We have observed that both generation time and biofilm formation capacity are significantly affected in the Δ*cntL* strain in comparison with the wild type (figure 1). We therefore suggest that the pseudopaline-dependent fitness phenotypes observed in MCM are also occurring during infection and therefore contribute to the pseudopaline requirement during infection. In addition to fitness deficiencies, we also observed a significant reduction in the global extracellular protease activity of the pseudopaline deficient strain (figure 1C). Taking into account that around 6% of bacterial proteomes includes Zn metalloenzymes (20, 21), involved in various cellular processes including translation, general metabolism or extracellular protease activities, we consider that the observed phenotypes are due to an impairment of Zn containing extracellular proteases. As a whole, these data indicate that several metalloenzymes important for infections rely on pseudopaline-mediated Zn uptake and justify the necessity of this metal uptake for *P. aeruginosa*’s fitness during lung infections occurring in cystic fibrosis patients. Therefore, any new additional information on the molecular mechanisms underlying this metal uptake pathway would be useful to understand it better and envisage its inactivation.

**Figure 1:**
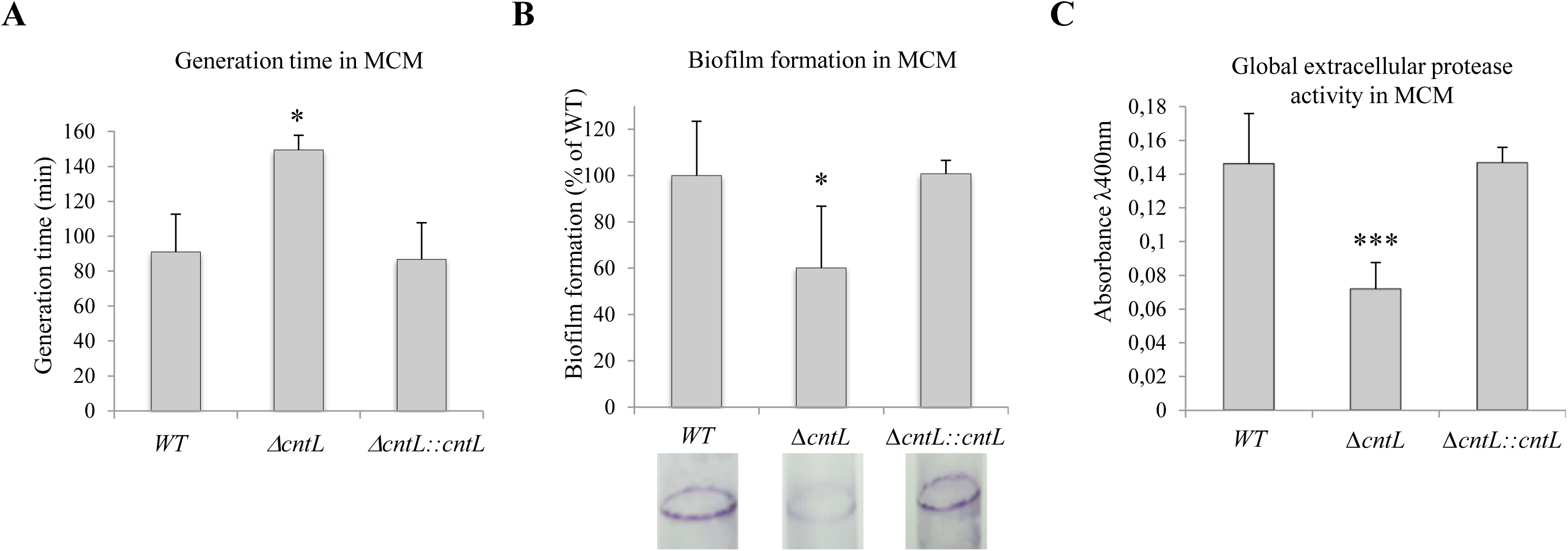
Impact of pseudopaline on generation time, biofilm formation capacity and extracellular protease activity in MCM. PA14 wild-type (WT), mutant (*ΔcntL)* and complemented (*ΔcntL::cntL*) strains were grown in MCM. Generation time (A), biofilm formation capacity (B) and extracellular protease activity (C) were measured as described in the M&M section. A representative illustration of crystal violet rings obtained with the different strains during the biofilm formation assay is presented panel B. Errors bar, mean ± standard deviation (sd) of at least two independent biological replicates. *P<0.05, ***P<0.001 as compared to the wild type (WT).

### Pseudopaline export across the bacterial OM is mediated by the MexAB-OprM efflux pump

We demonstrated elsewhere that pseudopaline was secreted in the medium during bacterial growth in MCM (7). While pseudopaline is exported across the IM by CntI, the machinery involved in its export across the OM remains unknown. In a preliminary search for pseudopaline OM export candidates, we selected four relevant RND efflux pumps of *P. aeruginosa* which are known to accommodate a wide array of structurally unrelated antibacterial molecules, namely MexAB-OprM, MexCD-OprJ, MexEF-OprN and MexXY/OprM (22). To test the possible involvement of the four selected efflux pumps in the pseudopaline secretion, we measured and compared the extracellular pseudopaline levels in a set of PA14 cultures including a quadruple mutant strain lacking the four MexAB, CD, EF and XY RND components, called PA14Δ*RNDs* and additional control strains deficient in pseudopaline synthesis (PA14Δ*cntL*) or IM export (PA14Δ*cntI*) (figure 2). The strains were grown in MCM and pseudopaline content was measured by hydrophilic interaction liquid chromatography (HILIC) – electrospray ionization mass spectrometry (ESI MS) in the culture supernatant. As already reported by Lhospice *et al*., (7), the PA14Δ*cntL* strain lacking the CntL enzyme shows no detectable pseudopaline in the supernatant while the PA14Δ*cntI* strain lacking the pseudopaline IM exporter CntI displays a substantial 90% decrease of the extracellular pseudopaline content (figure 2). Remarkably, the quadruple PA14Δ*RNDs* mutant strain displays a similar major reduction of the extracellular pseudopaline content, thus, indicating a similar extent of impairment in pseudopaline secretion in absence of the four RND efflux pumps. Therefore, we conclude that at least one of the four RND components selected is involved in pseudopaline translocation through the OM. We then focused our investigations on MexAB-OprM since it is the major and the most ubiquitous, constitutively expressed, RND efflux pump of *P. aeruginosa*. We first tested the involvement of the IM module of the pump, the MexAB RND component, by measuring extracellular pseudopaline content in the corresponding m*exAB* mutant strain (PA14Δ*mexAB*). This strain showed a similar reduced pseudopaline content to the PA14Δ*cntI* and PA14Δ*RNDs* strains, indicating that the MexAB RND component of the tripartite efflux pump MexAB-OprM is required for pseudopaline secretion. To confirm the involvement of this RND efflux pump in pseudopaline secretion, we tested the contribution of its OM TolC-like component, OprM to the extracellular pseudopaline recovery. Data presented figure 2 indicate that, in absence of OprM, pseudopaline secretion is impaired to the same extend than in the MexAB deficient strain. The similar impairment in pseudopaline secretion independently observed in strains lacking MexAB or OprM, known to form the tripartite MexAB-OprM RND efflux pump, revealed the exclusive involvement of this pump in pseudopaline secretion across the OM.

**Figure 2:**
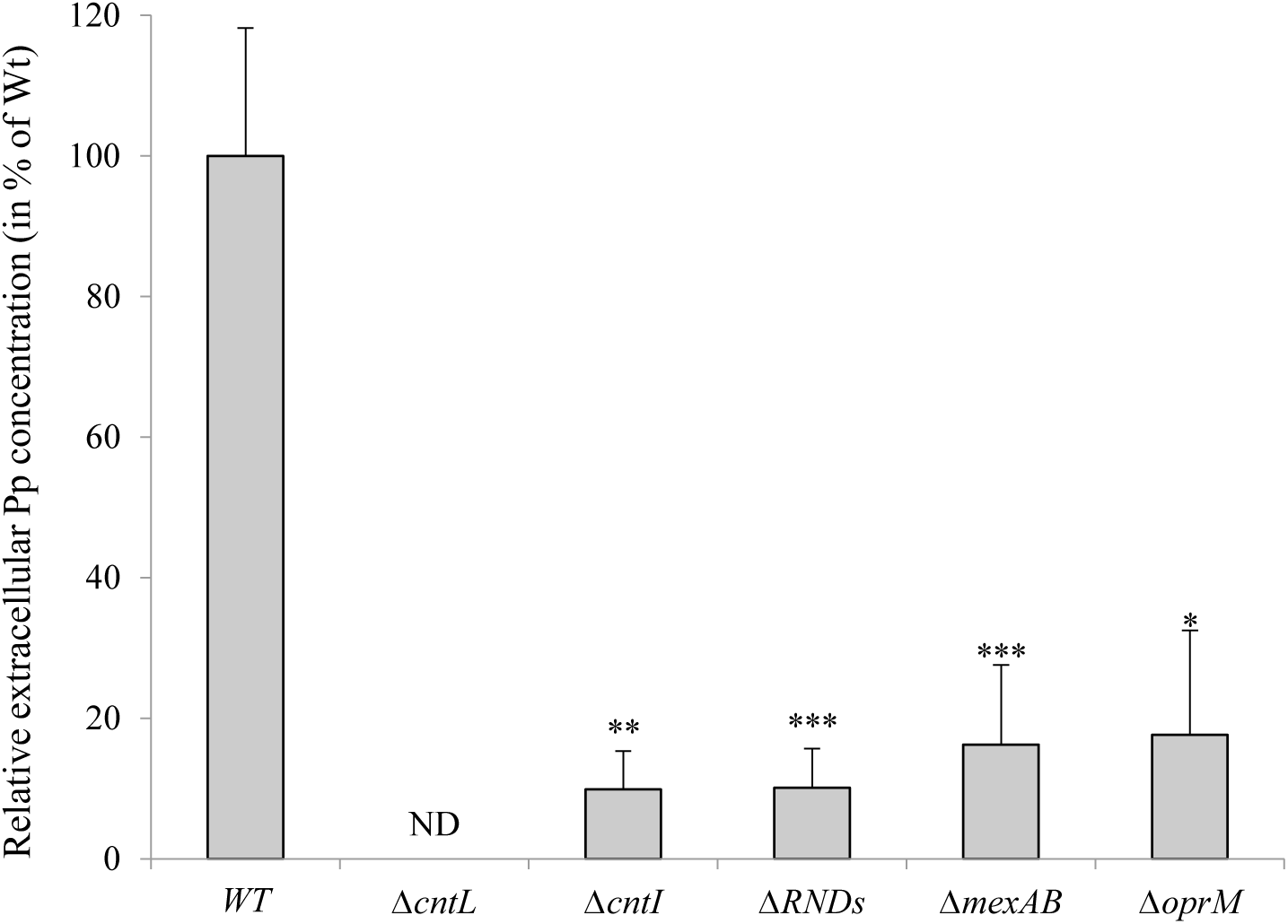
Extracellular pseudopaline content in absence of pseudopaline inner and outer membrane exporters. *P. aeruginosa* PA14 wild-type and mutant strains were grown for 10h at 37 °C in MCM. The relative levels of extracellular pseudopaline in the supernatant were measured by HILIC - ESI-MS. Errors bar, mean ± sd of at least three independent biological replicates. *P<0.05, **P<0.01, \*\*\**P*<0.001, as compared to the wild type (WT); ND: Not Detectable, NS: Not Significant, Pp: pseudopaline.

### The absence of pseudopaline OM exporter does not trigger intracellular pseudopaline accumulation

Pseudopaline is synthetized in the cytoplasm and then exported across the IM into the periplasm through CntI. Data presented in figure 2 show unambiguously that pseudopaline is taken in charge in the periplasm by the MexAB-OprM tripartite RND efflux pump to complete its secretion to the extracellular milieu, therefore constituting the second step of the secretion process. As already noticed earlier and further confirmed by this study, the intracellular pseudopaline measurement in absence of its IM exporter CntI revealed an important cytoplasmic accumulation, which is associated with a toxic phenotype (Figure 3). We showed that this phenotype was a consequence of pseudopaline accumulation since it is suppressed in the double *cntL*/*cntI* mutant which does not produce pseudopaline (figure 3B). In contrast, when pseudopaline OM export is impaired in strains lacking the MexAB or the OprM component of the MexAB-OprM RND efflux pump involved in this process, we do not detect any cellular accumulation (figure 3A) and did not observe any adverse effect on bacterial growth. (figure 3B). Consequently, the combined secretion defect of a MexAB-OprM deficient strain without periplasmic accumulation suggests that the periplasm is a compartment which cannot accommodate pseudopaline accumulation. Two hypotheses therefore arise: (i) either there is a regulative effect with a negative feedback on pseudopaline synthesis or (ii) it exists a specific periplasmic machinery preventing pseudopaline accumulation in this compartment.

**Figure 3:**
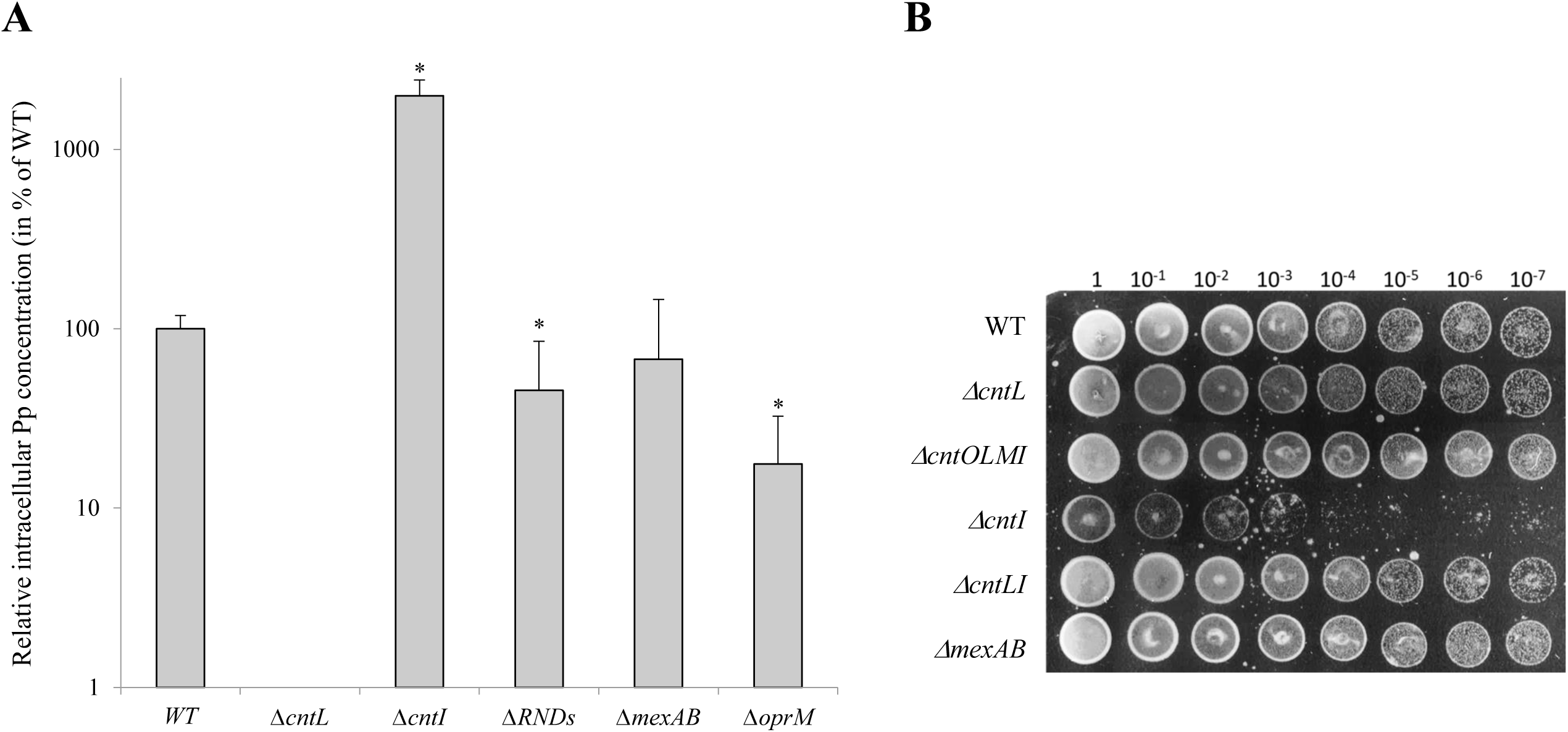
Intracellular pseudopaline content and phenotypes of *P. aeruginosa* PA14 strains lacking the pseudopaline inner and outer membrane exporters. (A) Detection of pseudopaline in the intracellular fractions of wild type and mutant strains. Strains were grown for 10h at 37 °C in MCM. The relative levels of intracellular pseudopaline in the cell fraction were measured by HILIC - ESI-MS. Errors bar, mean ± sd of at least three independent biological replicates. *P<0.05 as compared to the wild type (WT); ND: Not Detectable. (B) Viability of PA14 WT and mutant strains assessed by serial dilutions spotted on 1.5% agar MCM plates.

### Absence of pseudopaline OM exporter does not affect *cnt* gene’s expressions

The simultaneous decrease of the pseudopaline content in the supernatant of a PA14Δmex*AB* strain without a concomitant rise of pseudopaline’s periplasmic content (figures 2 & 3) raises the question of pseudopaline fate when it is not released in the milieu from the periplasm. In order to explore the hypothesis of a regulation mechanism of the *cnt* operon when RND efflux pumps are impaired, we examined the transcription level of the *cnt* operon in the different genetic backgrounds (figure 4). We used qRT-PCR to measure the transcriptional level of *cntO* and *cntM* genes in the PA14, PA14Δ*cntI*, PA14Δ*mexAB* and PA14Δ*RNDs* strains in MCM, a growth medium previously shown to induce pseudopaline production and secretion (figure S1). We found that the expressions of *cntO* and *cntM* genes of the *cnt* operon are not affected in the *cntI* mutant or in any RND efflux pump deletion combinations in comparison with the wild type. This allowed us to rule out the possibility of a negative feedback on the *cnt* genes expression when pseudopaline secretion is impaired. We therefore conclude that the pseudopaline production level is not affected by the deletion of the genes involved in its export across the bacterial envelope and the lack of extracellular pseudopaline observed in absence of MexAB-OprM is truly due to a secretion defect.

**Figure 4:**
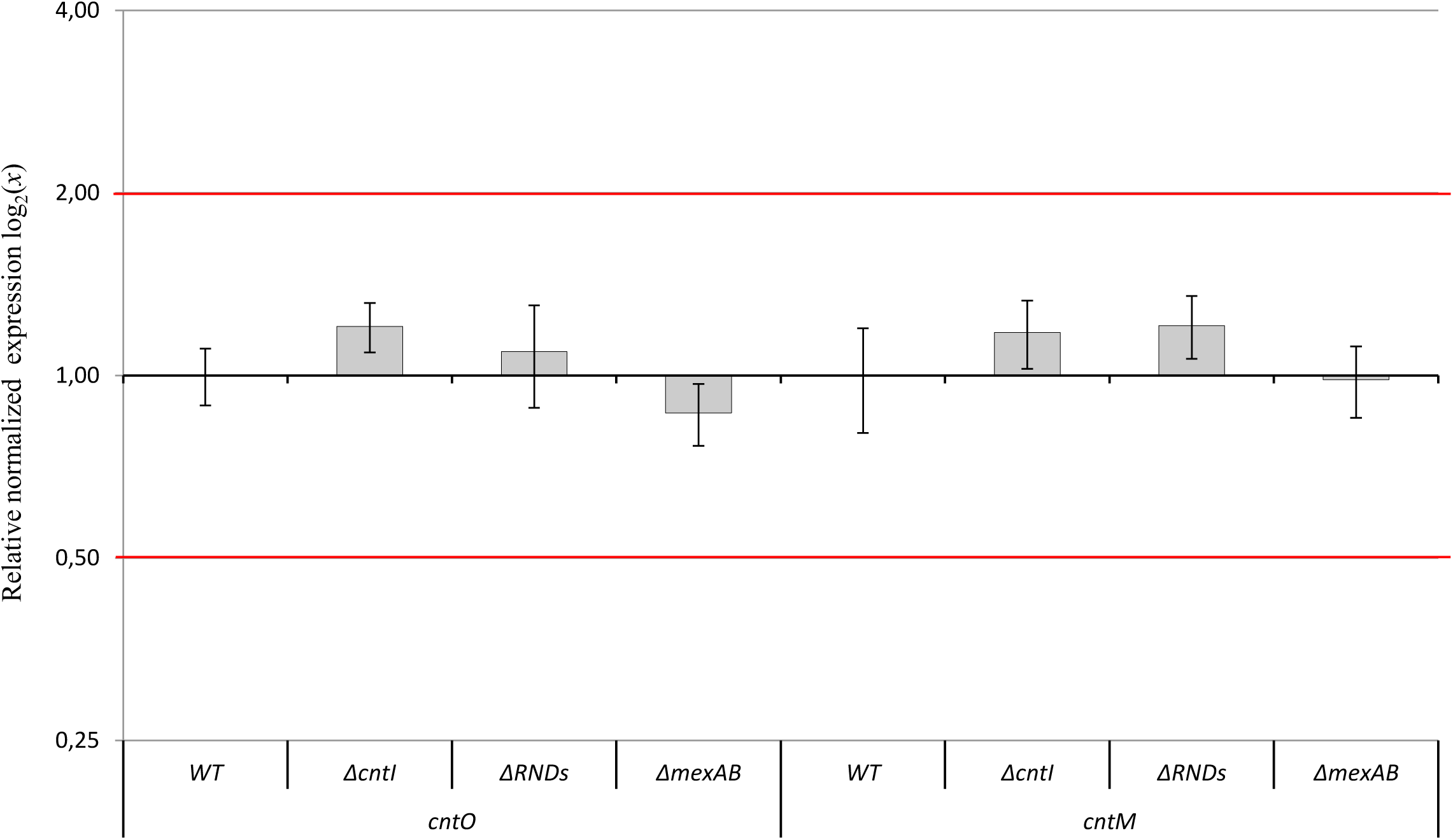
Expression of *cntO* and *cntM* genes in absence of pseudopaline inner and outer membrane exporters. Transcriptional activities were measured by RT-qPCR in PA14 wild type and various mutant strains. The relative *16S RNA* and *uvrD* normalized transcriptional expressions of *cntO* and *cntM* in various *cnt* and RNDs mutant strains compared to the wild type strain is presented. The two red lines are indicating the two time up and down regulation boundaries for a significant transcriptional effect. Experiments have been done in three biological replicates each including three technical replicates.

### Evidence of an intracellular pseudopaline modification mechanism

Since we could not detect periplasmic accumulation of pseudopaline when its export across the OM was impaired, we hypothesized that the non-secreted pseudopaline is either degraded or modified in the periplasm by a dedicated mechanism. To check this hypothesis, we took advantage of the reconstituted *in vitro* pseudopaline biosynthesis pathway previously developed (7) to follow the fate of radiolabeled pseudopaline under various conditions (figure 5). We used thin layer chromatography (TLC) to separate and visualize the radiolabeled pathway intermediate yNA and pseudopaline when the radiolabeled substrate ^14^C S-adenosyl-methionine (SAM) is incubated with defined substrate and enzymes (figure 5A top). We then followed by TLC the integrity of the *in vitro* synthesized radiolabeled pseudopaline upon incubation with various concentrations of PA14 cell extracts generated under *cnt* induction conditions. Data presented figure 5A reveal that the amount of radiolabeled pseudopaline gradually decreased upon increasing amount of PA14 lysates, with a doubling of the band corresponding to pseudopaline (figures 5A lines 4 to 6 and 5B lines 4 & 6), which reflects the appearance of a modified form of pseudopaline. This suggests the existence of a pseudopaline modification mechanism in the *P. aeruginosa* cellular soluble extracts. As a specificity control, no degradation was observed when *E. coli* cell lysates are added to the reaction in the same proportions (figures 5A lines 7 to 9 and 5B lines 8 & 9). Since pseudopaline accumulates in the cytoplasm in absence of IM exporter but does not accumulate in the periplasm in absence of OM exporter, our observation implies that the highlighted modification mechanism is restricted to the periplasmic compartment. Interestingly, the fact that pseudopaline modification is observed with both wild type PA14 and the PA14Δ*mexAB* lysates (figure 5B lines 4, 5 and 6, 7) suggests that this modification mechanism is constitutive rather specifically induced upon pseudopaline accumulation in absence of its OM exporter. We therefore propose that this pseudopaline modification process also occurs under physiological conditions and is used for metal recovery from the pseudopaline-metal complex imported *via* the CntO outer membrane importer (see model, figure 6).

**Figure 5:**
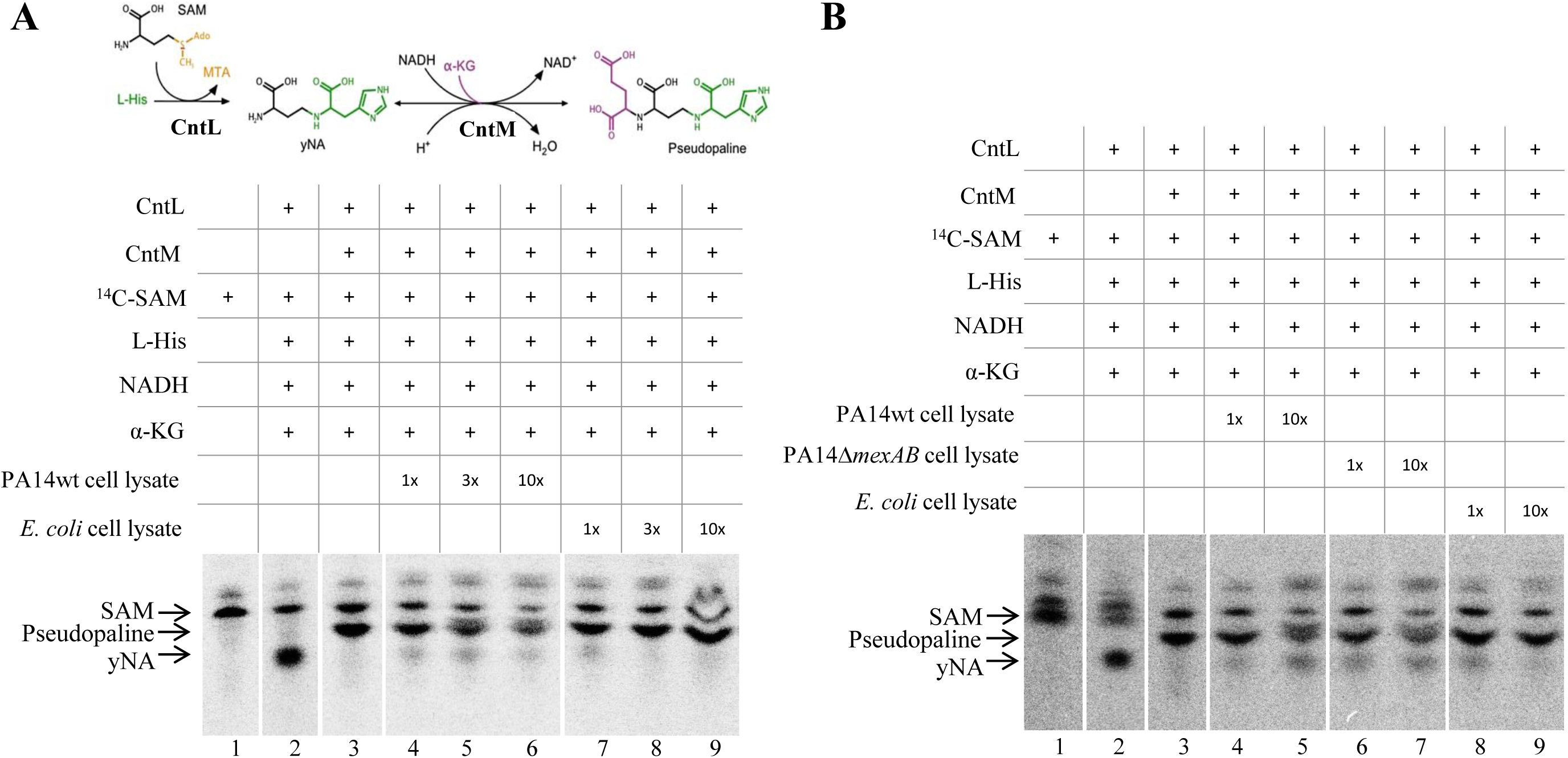
*in vitro* pseudopaline degradation by PA14 cell lysates. (A top) Summary of the CntL/M-dependent biosynthesis pathway for the assembly of yNA and pseudopaline from L-his, SAM, NADH and α-KG. (A, B) Thin layer chromatography (TLC) separation of [^14^C]-SAM (S-adenozyl methionine) and reaction products (yNA and pseudopaline) after incubation with defined enzymes, substrates and cofactor, followed or not by an incubation with increased concentration of *E. coli*, PA14wt and PA14Δ*mexAB* cell lysates. Incubation times for pseudopaline production are 15 and 10 minutes for experiments performed in panels A and B respectively. Incubation times with cell lysates are 15 and 45 minutes for experiments performed in panels A and B respectively.

**Figure 6:**
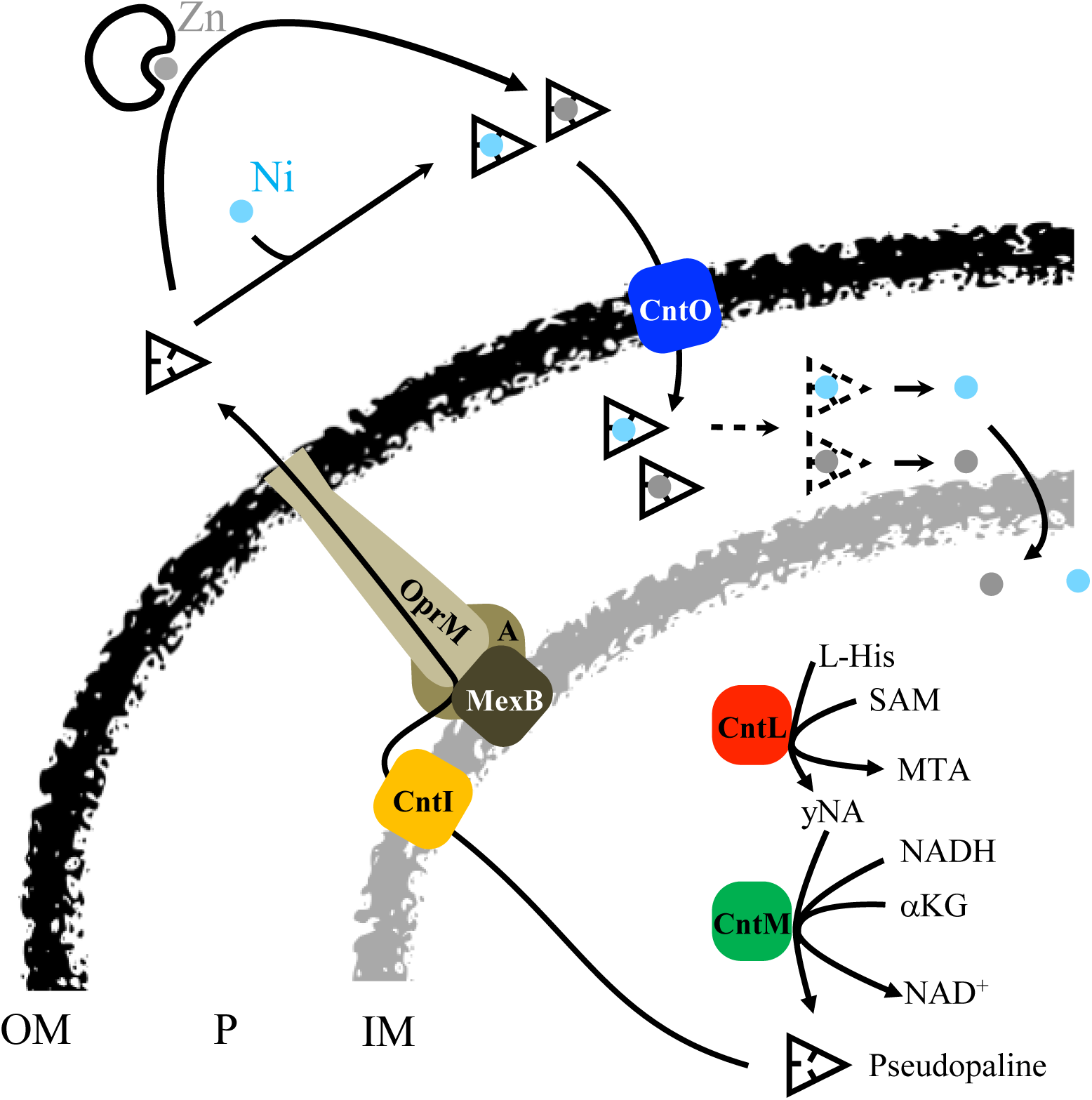
Model of pseudopaline synthesis, secretion, and metal uptake in *P. aeruginosa*. See text for details. Outer membrane (OM), periplasm (P), inner membrane (IM), L-histidine (L-His), S-adenozyl methionine (SAM), α-keto-glutarate (α-KG) and cofactor (NADH). The dashed triangle around pseudopaline symbolizes pseudopaline periplasmic degradation/modification.

### A model of pseudopaline secretion and recovery by *P. aeruginosa*

We described recently the metallophore pseudopaline, secreted by *P. aeruginosa* and involved in zinc uptake under metal limiting conditions (7). Owing to the data reported in this study on pseudopaline secretion and fate after recovery, we are now able to provide a detailed model of secretion and recovery of this important metallophore by *P. aeruginosa* (figure 6). In this updated model, following the synthesis of pseudopaline in the cytoplasm by the CntL and CntM enzymes, the metallophore is then translocated into the periplasm through the CntI exporter. As demonstrated in this study, the periplasmic pseudopaline uses the MexAB-OprM RND efflux pump for the final release in the extracellular milieu. MexAB-OprM therefore constitutes, together with CntI, a two-step secretion system that exports pseudopaline from the site of its synthesis across the IM, periplasm and OM. Our data reveal the involvement of the MexAB-OprM efflux pump in the secretion of a natural component made by the bacteria that may constitute one of its primary substrates. In the extracellular medium, pseudopaline binds Zn and Ni. It is then recognized by the OM TonB-dependent receptor CntO which allows its import into the periplasm. Additional results generated in this study strongly suggest the existence of a pseudopaline modification system involved in periplasmic pseudopaline homeostasis and possibly in metal liberation. The free metal will then be available to bacterial metalloproteins following its import into the cytoplasm by general importers such as ZnuA in the case of zinc (11)

## DISCUSSION

Pseudopaline is a metallophore synthetized and used by the opportunistic bacterial pathogen *P. aeruginosa* to recover metals from metal scarce environments. This metallophore plays an important role in bacterial survival during infection likely resulting from the lack of bio-available metals in the host. The pseudopaline metal uptake pathway utilizes a complex secretion and recovery cycle of the metallophore across the Gram-negative bacterial envelope. We showed previously that pseudopaline export across the inner membrane and import across the outer membrane are mediated by the CntI exporter and the CntO TBDT, respectively, both encoded on the pseudopaline *cnt* operon together with the *cntL* and *cntM* biosynthetic genes. In this study, we are extending our previous characterization of the pseudopaline cycle by revealing the involvement of an RND efflux pump in this process. Our data demonstrate that the MexAB-OprM efflux pump is responsible for the final secretion step of the metallophore in the extracellular medium therefore filling an important gap in the pseudopaline cycle.

In the molecular mechanisms underlying pseudopaline secretion, MexAB-OprM forms together with CntI an unprecedented two-step secretion system that drives pseudopaline from the cytoplasm to the extracellular medium. The involvement of an efflux pump in the cellular release of a metallophore has already been reported but in combination with different IM exporters. The *E*.*coli* enterobactin is exported across the cytoplasmic membrane through the IM major-facilitator transporter EntS (23) and then exported across the OM from the periplasm through the OM channel TolC, an ortholog of the *P. aeruginosa* OprM (19). Another example was reported in the closely related organism *Salmonella enterica* Sv Typhimurium where the salmochelin is exported to the medium through the IroC-TolC complex (24). This is also the case in *P. aeruginosa* where the siderophore pyoverdine is secreted through the PvdRT-OpmQ efflux pump (17, 18). Here we unravel an unprecedented combination involving an IM exporter belonging to the EamA or drug/metabolite transporter (DMT) family and the MexAB-OprM RND efflux pump and, therefore, extend the spectrum of efflux pump-associated two-step secretion systems involved in metallophores across Gram-negative envelopes. On the basis of structural and functional studies, it was previously described that the RND protein AcrB, a close homolog of the MexB protein, is a homotrimer acting as a peristaltic pump, able to load its substrate in the periplasmic compartment before secretion across the OM through the TolC channel (25). However, whether efflux pump-dependent metallophores including pseudopaline are first exported to the periplasm, remains to be determined.

So far, MexAB-OprM efflux functions have mostly been dedicated to the release of exogenous toxic compounds such as antibiotics as well as a series of amphiphilic molecules, disinfectants, dyes, solvents or detergents (26). Additional data indicate that MexAB-OprM efflux pump is also releasing non-permeable autoinducers involved in bacterial quorum sensing (27, 28). The present study highlights an additional function for the MexAB-OprM efflux pump in endogenous metallophore secretion. Such involvement of MexAB-OprM in the secretion of important small natural compounds has already been indirectly suggested following the observation that a *mexAB* mutant is unable to kill mice that were deficient in leucocytes, whereas the parent strain caused a fatal infection (29). The finding that MexAB-OprM is essential for pseudopaline secretion, itself necessary to support *P. aeruginosa* growth in airway mucus secretions (4) provide a possible additional explanation for the MexAB-OprM requirement in virulence.

While at least 12 different functional RND efflux pumps have been described in *P. aeruginosa*, our data reveal a specific and exclusive recognition of pseudopaline by MexAB-OprM. In a recent study, Tsutsumi *et al*. revealed the structure of the complete MexAB-OprM complex in the presence or absence of drugs at near-atomic resolution and proposed precise mechanisms for substrate recognition and efflux supporting such specificity (25). In addition, Ramaswany *et al*. recently performed an exhaustive atomic-level comparison of the main putative effector recognition sites in the two RND modules MexB and MexY (30). They pointed out specific signatures that may constitute selectivity filters such as mosaic-like lipophilic and electrostatic surfaces of the binding pockets of MexB and MexY proteins providing multifunctional binding sites for diverse substrates. Further structure function investigations are necessary to understand MexB specificity toward pseudopaline. The question, whether the natural endogenous compound pseudopaline is recognized and exported in a same manner as the exogenous compounds requires further investigations.

In contrast to pyoverdine (17, 18) and enterobactin (31), we observe neither subsequent accumulation, nor toxicity when pseudopaline export across the OM is impaired. We are providing experimental data suggesting that the absence of pseudopaline accumulation in these conditions is due to pseudopaline modification by a dedicated system specific to *P. aeruginosa*. The presence and activity of such a pseudopaline modification mechanism in other than abnormal pseudopaline periplasmic accumulation, suggests a function for this machinery under physiological conditions in the wild type bacteria. The absence of encoded specific machinery in the *cnt* operon to import the pseudopaline-metal complex through the IM into the cytoplasm (as found for example in the *cnt* operon in *S. aureus* and *Y. pestis* (8)) is in line with the hypothesis of metal release from pseudopaline in the periplasm. We therefore propose that the pseudopaline-mediated metal import pathway ends in the periplasm, where the free metal will be distributed to various metalloproteases such as exoproteases that fold in the periplasm before secretion in the medium by the type II secretion system (32) or cytoplasmic metalloenzymes after metal transport through pseudopaline-independent importers such as Znu ABC transporter described for zinc import (11). Further experiments, out of the scope of this study, will be necessary to identify the determinants underlying the pseudopaline’s modification mechanism.

In the case of pyoverdine, it has been shown that it can bind and import in the periplasm a large array of metals with different affinities, (33). In the same study, the authors demonstrated that the PvdRT-OpmQ machinery is involved in the secretion of unwanted siderophore-metal complexes, leading to specific integration of iron in the cellular pool. Therefore, we hypothesize that, similarly, MexAB-OprM could play a role in the selectivity of metals imported by pseudopaline. Investigation on this potential role would give an insight on the nature of metal physiologically targeted by this machinery. Indeed, if the import of zinc by pseudopaline is required for full extracellular metalloprotease activity (Figure 1), the physiological relevance underlying Ni and Co import by pseudopaline remains to be demonstrated.

Finally, other siderophores such as pyoverdine have been shown to be recycled (34), and this could also be the case for pseudopaline. The absence of a recycling mechanism would lead to a constant production of pseudopaline during infection, and would represent an important investment of cellular resources. More research is needed in order to determine whether the modification of pseudopaline we observed is reversible and allows recycling or, alternatively, whether pseudopaline is fully degraded after uptake.

## MATERIAL AND METHODS

### Bacterial strains, plasmids and liquid growth conditions

Bacterial strains, vectors and plasmids used in this study are listed in Table 1. *E. coli* strains were grown aerobically with orbital shaking at 37 °C in Luria-Broth (LB) with antibiotics as required (25 μg ml^−1^ kanamycin (Kan), 25 μg ml^−1^ tetracycline (Tc), 15 μg ml^−1^ gentamicin (Gm), 30 μg ml^−1^ streptomycin (Sm)). The *E. coli* strains CC118λpir and SM10 were used to propagate pKNG101 derivatives mutator plasmids. Recombinant plasmids were introduced into *P. aeruginosa* by triparental mating using pRK2013 and transconjugants selected on *Pseudomonas* isolation agar (PIA, Difco Laboratories) supplemented with antibiotics as required (500 μg ml^−1^ carbenicillin (Cb), 150 μg ml^−1^ Gm, 2000 μg ml^−1^ Sm, 200 μg ml^−1^ Tc). All *P. aeruginosa* strains used in this study were derivatives of the parental PA14 strain. *P. aeruginosa* strains were grown aerobically with horizontal shaking at 37 °C with antibiotics as required (150 μg ml^−1^ (Cb), 50 μg ml^−1^ (Gm), 500 μg ml^−1^ (Sm), 50 μg ml^−1^ (Tc)). Growths were performed rich LB medium and in minimal chelated medium (minimal succinate media (per liter: 30g K_2_HPO_4_, 15g KH_2_PO_4_, 5g (NH_4_)_2_SO_4_, 1g MgSO_4_, 20g Succinic acid and 15.5g NaOH, pH 7.0, MS) supplemented with 100µM EDTA, MCM). MCM pre-culture of 20 mL were inoculated from fresh MCM, 1.5% agar plates, grown until late exponential phase in polycarbonate Erlenmeyer flasks in order to avoid any metal contamination. Growth was monitored by OD_600_ measurement.

**Table 1:**
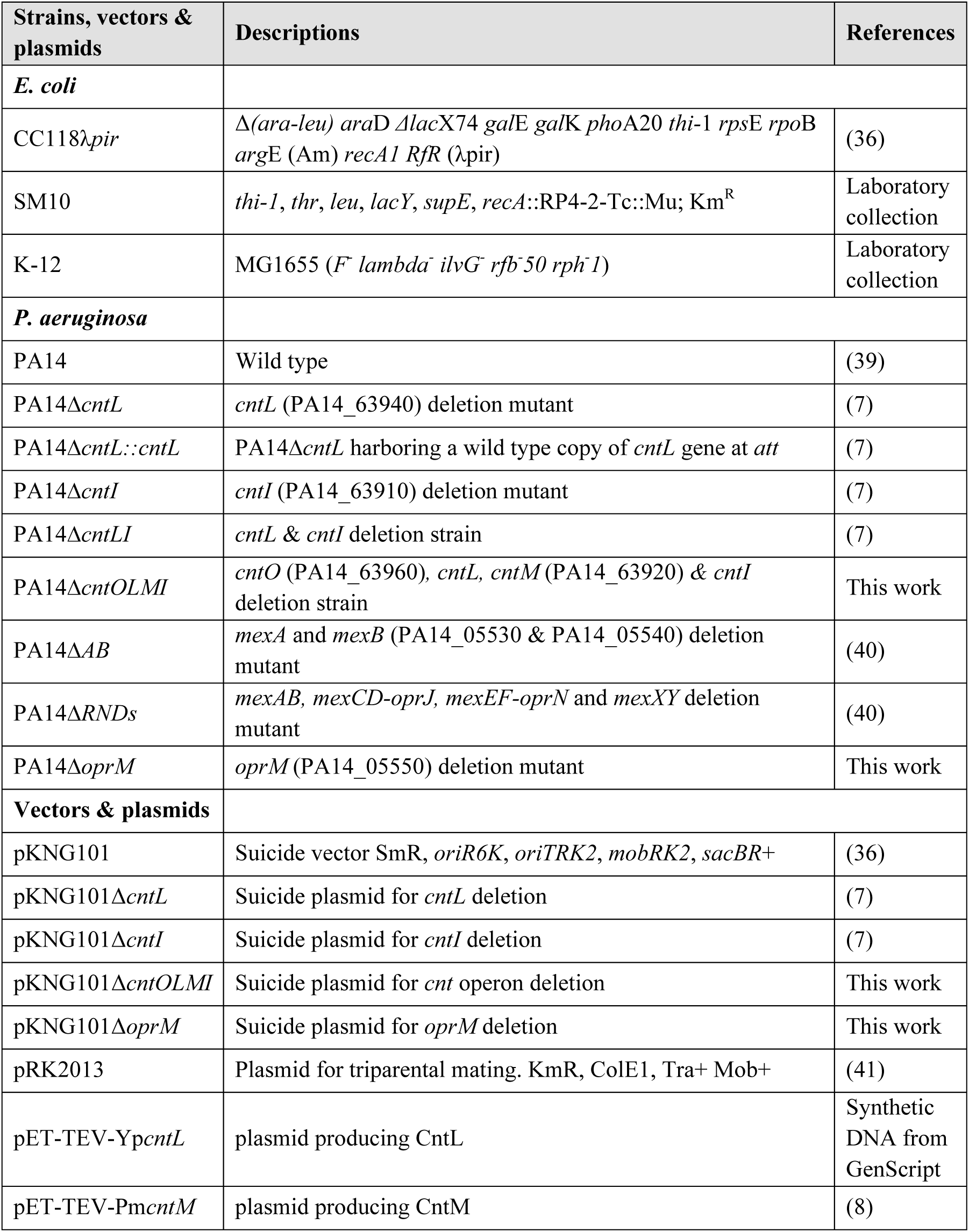
Bacterial strains, vectors and plasmids used in this study.

### Plasmid constructions

All plasmids constructed in this study except pKNGΔ*oprM* were generated by the one-step sequence and ligation-independent cloning (SLIC) method described in (35). To construct the suicide plasmid pKNGΔ*oprM*, the Δ*oprM* fragment was cloned into plasmid pCR-Blunt according to manufacturer’s instructions (invitrogen) and then subcloned as BamHI/XbaI fragment into the suicide vector pKNG-101. All PCR primers employed for plasmid construction are listed in Table 2. Genomic DNA was isolated and purified with Pure Link genomic DNA minikit (Invitrogen). PCR reactions for cloning were performed by using Q5® High-Fidelity DNA Polymerase (New England Biolabs, Inc (NEB). All the products sequenced to verify the absence of any mutations (GATC-biotech).

**Table 2:**
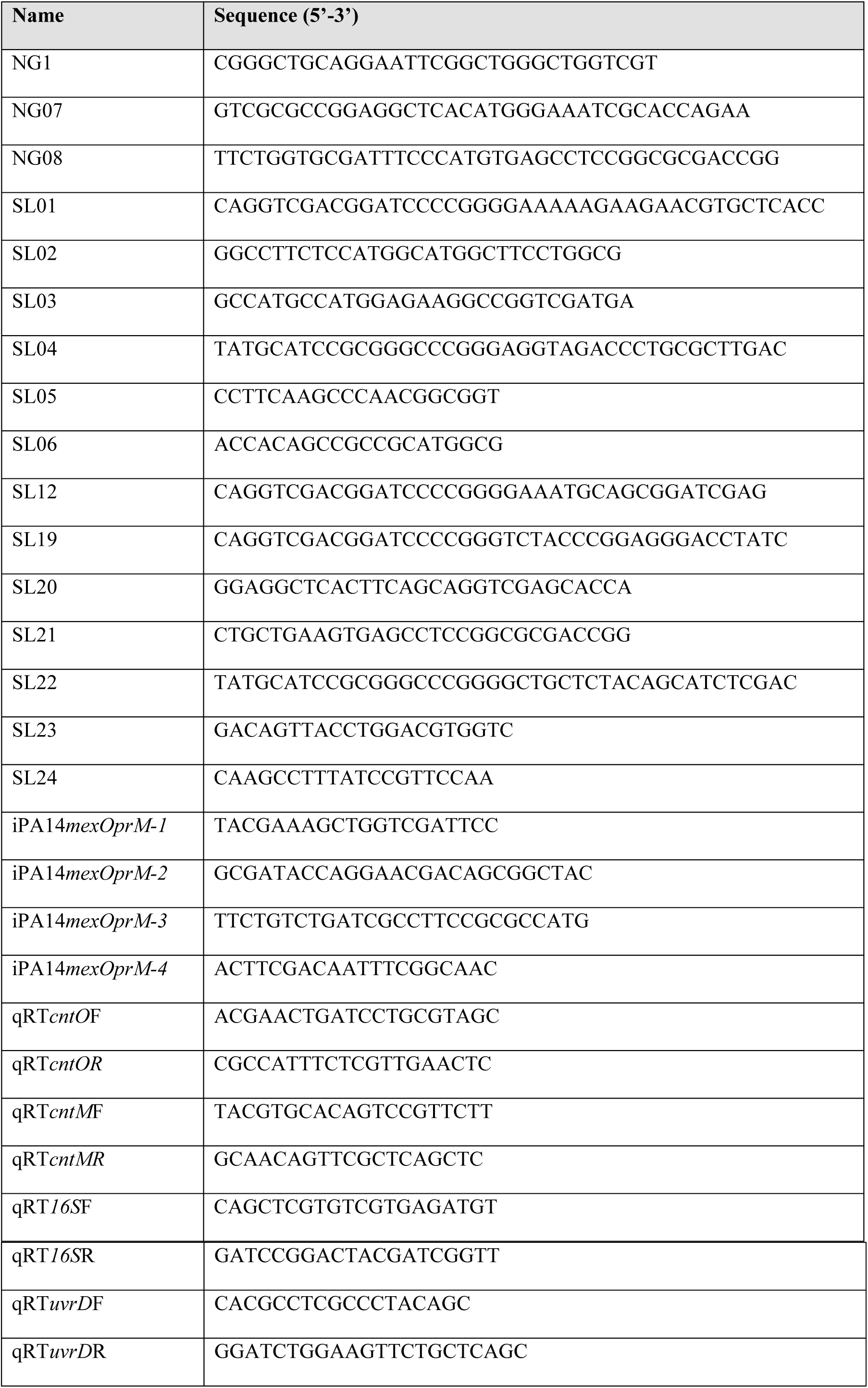
Oligonucleotides used in this study.

### Construction of *P. aeruginosa* PA14 knock-out mutants

PA14Δ*oprM* was constructed from PA14 using overlapping PCRs with specific iPA14*mexOprM* primers described in Table 2. Homologous recombination was carried out between the 5’ (512-bp) and 3’ (539-pb) regions flanking *oprM* according to the protocol of (36) modified by (37). The allelic exchange was verified by PCR and nucleotides sequencing experiments confirmed the deletion of 1415-bp in the *oprM* locus. To construct the pKNG101Δcnt*OLMI*, two DNA fragments corresponding to upstream and downstream regions of *cntO* and *cntI* were amplified from PA14 chromosomal DNA with PCR primers SL12/NG07 and NG08/SL22, respectively Upstream (639-bp) and downstream (517-bp) regions were ligated by overlapping PCR and cloned into linearized pKNG101 by the SLIC method. The resulting constructs were transformed into *E. coli* CC118λpir and introduced into *P. aeruginosa* PA14 by conjugation. The strains in which the chromosomal integration event occurred were selected on *Pseudomonas* isolation agar Gm plates. Excision of the plasmid, resulting in the deletion of the chromosomal target gene was performed after selection on Luria-Bertani (LB) plates containing 6% sucrose. Clones that became sucrose resistant and Sm sensitive were confirmed to be deleted for the genes of interest by PCR analysis.

### Generation time, biofilm formation and extracellular protease activity measurements

Generation time in minutes defines the time required by the bacterium to perform one cycle of division. Generation time was calculated in exponential phase from the growth curves of six independent biological replicates. Each over-night cultures were inoculated at OD_600_ of 0,05 in fresh MCM in clear, flat bottom, 96-well plates (Corning™ Costar 3596). Growth was recorded in a Tecan Spark Microplate Incubator/Reader (Thermo Fisher Scientific) for 24 hours at 37°C under continuous shaking and with 600nm absorbance readings every 30 minutes, corrected with the absorbance of non-inoculated media.

Biofilm formation was measured in clear, flat bottom, 24-well plates (Corning™ Costar 3526). Each over-night cultures were inoculated at OD_600_ of 0,2 in 1 mL fresh MCM and incubated at 30°C for 24 hours. Following incubation, planktonic bacteria and media were rinsed away with non-sterile deionized water. The wells were filled with 0,1% crystal violet solution and sat for 15min at RT and then washed with deionized water three times. Any crystal violet staining on the bottom of the well were cleaned, and the plate was let to air-dry overnight at room temperature. Crystal violet rings were then solubilized in 30% glacial acetic acid. Adherent biofilm was quantified by measuring optical density of all samples at 550nm and normalized to WT mean absorbance. Photos used for biofilm illustration where made from biofilm experiments following the same method, but realized in borosilicate glass tubes for microbiological cultures.

Global extracellular protease activity: Pre-culture of PA14 strains are inoculated from a fresh 1,5% agar MCM plate, and incubated overday in 20 mL MCM under horizontal shaking at 37°C. A 25 mL culture of fresh MCM is then inoculated by the pre-culture at OD_600_ of 0,05 and incubated for 10 hours under horizontal shaking at 37 °C in polycarbonate baffled Erlenmeyer flask (Corning 431407). Culture were centrifuged at 13000 x*g* for 15 min at 4°C and supernatants was concentrated with Amicon Ultra-15 10000Da NMWL (Merk C7715) and normalized to cell density. Total extracellular proteolytic activity was determined using casein conjugated to an azo-dye as a substrate, following a modified procedure described by Kessler *et al*., (38). Briefly, 0,25 mL of concentrated and normalized supernatant was added to 0,75 mL reaction buffer (0,05M Tris-HCl, 0,5 mM CaCl_2_, pH 7,5, 3 mg.mL^-1^ Azocasein (Sigma A2765)) preheated at 37°C and vortexed. The reaction mixture was incubated at 37°C for 15 min. The indigested substrate was precipitated by adding trichloroacetic acid to a final 3,3% and vortexed. The samples were then left at RT for 30 minutes, and centrifuged at 10000 x*g* for 20min. The absorbance was measured at 400 nm.

### Viability assay on plate

Pre-culture of PA14 strains were performed overnight in MCM under horizontal shaking at 37°C. The next day, a culture of fresh MCM is inoculated by the pre-culture at OD_600_ of 0.05 and incubated for 9 hours under horizontal shaking at 37 °C. Cultures were then adjusted to OD_600_ of 1 and subjected to 10% serial dilutions in fresh MCM. 10 μl of culture samples were spotted on MS 1.5% agar or agarose plate an incubated 30h at 37 °C.

### RNA Preparation and Reverse Transcription

RNAs were prepared from 25 mL PA14 culture in MCM in the exponential growth phase after 10h shaking at 37°C. The cells were harvested and frozen at −80°C. Total RNAs were isolated from the pellet using RNeasy mini Kit (Qiagen) according to the manufacturer’s instructions; an extra TURBO DNAse (Invitrogen) digestion step was done to eliminate the contaminating DNA. The RNA quality was assessed by the TapeStation 4200 system (Agilent). RNA was quantified spectrophotometrically at 260 nm (NanoDrop 1000; Thermo Fisher Scientific). For cDNA synthesis, 1 µg total RNA and 0.5 μg random primers (Promega) were used with the GoScript™ Reverse transcriptase (Promega) according to the manufacturer’s instruction.

### Quantitative real-time PCR (qPCR)

qPCR analyses were performed on a CFX96 Real-Time System (Bio-Rad, France). The reaction volume was 15 μL and the final concentration of each primer was 0.5 μM. The cycling parameters of the qRT-PCR were 98°C for 2 min, followed by 45 cycles of 98°C for 5 s, 60°C for 10 s. A final melting curve from 65°C to 95°C was added to check the specificity of the amplification. To determine the amplification kinetics of each product, the fluorescence derived from the incorporation of EvaGreen into the double-stranded PCR products was measured at the end of each cycle using the SsoFast EvaGreen Supermix 2X Kit (Bio-Rad). The results were analyzed using Bio-Rad CFX Manager software, version 3.1 (Bio-Rad). The 16S RNA and *uvrD* genes were used as a reference for normalizations. For each point a technical duplicate was performed. The amplification efficiencies for each primer pairs were comprised between 80 and 100%. All of the primer pairs used for qPCR are shown in the table 2.

### Pseudopaline detection by HILIC - ESI-MS

All the *P. aeruginosa* strains were grown aerobically with horizontal shaking at 37 °C in freshly made MCM. Cultures were then adjusted at OD = 0.05 in 25 mL freshly made MCM and grown for 10 h. After 10 h, OD_600_ was measured before cells were harvested by centrifugation (4,000xg, 30 min, 4 °C). The supernatant was collected, filtered and stored at −80 °C. The pellet was washed three times with cold MCM and resuspended in 1.3 ml cold MCM. OD_600_ were measured, and cells ruptured by successive sonication cycles (three times 30 sec at 60% of maximal power, keeping the solution cold (Brandson Sonifier 450 microtip)). The lysates were then centrifuged at 16,000xg for 30 min at 4 °C and supernatants were collected, filtered at 0.22 µm, and stored at −80 °C. These cell lysate and supernatant fractions were analyzed to detect pseudopaline using HILIC/ESI-MS as previously described (7). Pseudopaline levels were determined using the relative intensity of the m/z corresponding to the pseudopaline-Ni complex in extracellular or intracellular fractions saturated with nickel, in order to force nickel complex formation.

### *in vitro* pseudopaline modification monitoring by thin layer chromatography

Preparation of *P. aeruginosa* and *E. coli* cell lysates. Pre-culture of PA14 strains are inoculated from a fresh MS + 1,5% agar plate, and incubated overday in 20 mL MS medium under horizontal shaking at 37 °C. The culture is then adjusted at OD_600_ = 0,05 with 100 mL of fresh MCM and incubated for 10 hours under horizontal shaking at 37 °C. *E. coli* MG1655 strain were grown in M63 media (per liter of water, 13.6 g KH_2_PO_4_, 2 g (NH_4_)_2_SO_4_, 0.2 g MgSO_4_, glucose 0.2%, casamino acids 0.5%, pH 7), with the same culture time and conditions. An aliquot corresponding to 100 OD_600_ units of each bacterial culture was centrifuged for 10 min at 2,000x*g*. Cell pellets were resuspended in 10 mL 50 mM Tris-HCl pH 7.6 and ruptured by successive sonication cycles (three times 30 sec at 60% of maximal power, keeping the solution cold (Brandson Sonifier 450 microtip). The final lysate was centrifuged at 2,000x*g* for 15 min to remove unbroken cells and the supernatant was saved as total cell lysate fraction.

Radiolabeled yNA and pseudopaline are first produced with the reaction of the purified enzymes CntL (for yNA) or a mixture of CntL and CntM (for pseudopaline) with their substrates and then incubated with cell lysates. For the production reaction, we used CntL from *Yersinia pestis* (YpCntL) and CntM from *Paenibacillus mucillaginosus* (PmCntM) that have been shown to be more stable and effective than their equivalent from *Pseudomonas aeruginosa* (8). Cloning, expression and purification of PmCntM have already been described (8). For YpCntL, the gene encoding the YpCntL protein was cloned in a pET-TEV plasmid (synthetic DNA from GenScript). After transformation, *E. coli* BL21 strains are aerobically cultivated with horizontal shaking in LB media supplemented with kanamycin at 50 mg.ml^−1^. These strains were grown at 37°C and, when the OD of the culture was ∼0.6, IPTG was added at a final concentration of 0.1 mM for induction. After overnight growth at 16°C, cells were recovered by centrifugation at 5,000xg for 20 min at 4°C, resuspended in buffer (20 mM Na_2_HPO_4_, 300 mM NaCl, 15 mM Imidazole, pH 7) and disrupted using a Constant cell disruption system operating at 2 Kbar. Cell debris were removed by centrifugation at 8,000xg for 20 min and the supernatants were centrifuged at 100,000xg for 45 min at 4°C to remove cell wall debris and membrane proteins. The resulting soluble fraction was loaded on a nickel affinity column (HisTrap 1 ml column, G.E. Healthcare), and the protein was eluted stepwise with imidazole (50 mM for wash and 500 mM for elution). Elution fraction was adjusted to 1 mg/mL and cleaved by TEV protease in a ratio 1:80 w/w. After an overnight incubation, the mixture was centrifuged at 10,000xg for 10 min and then loaded again on a nickel affinity column. YpCntL was recovered in the flow throw and then transferred into an imidazole-free buffer (20 mM Na_2_HPO_4_, 300 mM NaCl, glycerol 10%, pH 7).

For the TLC assays, the purified enzymes are incubated at a final concentration of 2.5 μM with carboxyl-[^14^C]-labeled SAM (2.5 μM), L-histidine (10 μM), NADPH (30 μM) and α-KG (1 mM). The total volume was 100 µl in ammonium acetate buffer (100 mM, pH=7). The mixtures are incubated for 10 or 15 min at 28°C. Cell lysates at different concentrations are then added to the radiolabeled pseudopaline (50% of each, v/v), and these mixtures are incubated for another 15 or 45 min at 28 °C. A same reaction is done with the addition of only ammonium acetate buffer in the radiolabeled pseudopaline as a control. The reactions are finally stopped by adding ethanol to a final concentration of 50% (v/v). An aliquot of 10 µl of the reaction mixtures is spotted on HPTLC (High-Performance TLC) Silica Gel 60 Glass Plates (Merck KGaA) after centrifugation at 16.000 g for 2 min. The products are then separated by thin-layer chromatography by developing plate with a phenol:n-butanol:formate:water (12 : 3 : 2 : 3 v/v) solvent system. HPTLC plates were dried and exposed to a [^14^C]-sensitive imaging plate for 5 or 6 days. Imaging plates are then scanned on a Typhoon FLA 7000 phosphorimager (GE Healthcare). The original TLC plates of *in vitro* pseudopaline degradation by PA14 cell lysates experiments presented in figure 4 are presented figure S2.

### Statistics

Statistics were determined using the Student’s t-test function of excel using a bilateral model and assuming equal variance when distribution of samples had normal distribution (figure 1). For nonparametric data (figures 2 and 3) statistical significance was calculated using the Wilcoxon rank sum test.

## FUNDING INFORMATION

This work was supported by the grant 20160501495 from “Vaincre la Mucoviscidose” and “Gregory Lemarchal” associations allocated to RV, PA and PP.

### ACKNOWLEDGEMENTS

We gratefully thank V. Pelicic, S. Lory, I. Broutin and J.F. Collet for careful reading of the Manuscript. We also thank the whole RV group for constant support in the project.

## CONFLICTS OF INTEREST

We have no conflict of interest

## Notes

### Competing Interest Statement

The authors have declared no competing interest.

## REFERENCES

1. Hood MI, Skaar EP. 2012. Nutritional immunity: transition metals at the pathogen-host interface. Nat Rev Microbiol 10:525–37.

2. Weinberg ED. 1975. Nutritional immunity. Host’s attempt to withold iron from microbial invaders. JAMA 231:39–41.

3. Johnstone TC, Nolan EM. 2015. Beyond iron: non-classical biological functions of bacterial siderophores. Dalton Trans 44:6320–39.

4. Gi M, Lee KM, Kim SC, Yoon JH, Yoon SS, Choi JY. 2015. A novel siderophore system is essential for the growth of *Pseudomonas aeruginosa* in airway mucus. Sci Rep 5:14644.

5. Nguyen AT, Oglesby-Sherrouse AG. 2015. Spoils of war: iron at the crux of clinical and ecological fitness of *Pseudomonas aeruginosa*. Biometals 28:433–43.

6. Schalk IJ, Cunrath O. 2016. An overview of the biological metal uptake pathways in *Pseudomonas aeruginosa*. Environ Microbiol 18:3227–3246.

7. Lhospice S, Gomez NO, Ouerdane L, Brutesco C, Ghssein G, Hajjar C, Liratni A, Wang S, Richaud P, Bleves S, Ball G, Borezee-Durant E, Lobinski R, Pignol D, Arnoux P, Voulhoux R. 2017. *Pseudomonas aeruginosa* zinc uptake in chelating environment is primarily mediated by the metallophore pseudopaline. Sci Rep 7:17132.

8. Laffont C, Brutesco C, Hajjar C, Cullia G, Fanelli R, Ouerdane L, Cavelier F, Arnoux P. 2019. Simple rules govern the diversity of bacterial nicotianamine-like metallophores. Biochem J 15:2221–2233.

9. Ghssein G, Brutesco C, Ouerdane L, Fojcik C, Izaute A, Wang S, Hajjar C, Lobinski R, Lemaire D, Richaud P, Voulhoux R, Espaillat A, Cava F, Pignol D, Borezee-Durant E, Arnoux P. 2016. Biosynthesis of a broad-spectrum nicotianamine-like metallophore in *Staphylococcus aureus*. Science 352:1105–9.

10. McFarlane JS, Davis CL, Lamb AL. 2018. Staphylopine, pseudopaline, and yersinopine dehydrogenases: A structural and kinetic analysis of a new functional class of opine dehydrogenase. J Biol Chem 293:8009–8019.

11. Pederick VG, Eijkelkamp BA, Begg SL, Ween MP, McAllister LJ, Paton JC, McDevitt CA. 2015. ZnuA and zinc homeostasis in Pseudomonas aeruginosa. Sci Rep 5:13139.

12. Mastropasqua MC, D’Orazio M, Cerasi M, Pacello F, Gismondi A, Canini A, Canuti L, Consalvo A, Ciavardelli D, Chirullo B, Pasquali P, Battistoni A. 2017. Growth of *Pseudomonas aeruginosa* in zinc poor environments is promoted by a nicotianamine-related metallophore. Mol Microbiol doi: 10.1111/mmi.13834.

13. Zhang J, Zhao T, Yang R, Siridechakorn I, Wang S, Guo Q, Bai Y, Shen HC, Lei X. 2019. De novo synthesis, structural assignment and biological evaluation of pseudopaline, a metallophore produced by *Pseudomonas aeruginosa*. Chem Sci 10:6635–6641.

14. Palmer KL, Mashburn LM, Singh PK, Whiteley M. 2005. Cystic fibrosis sputum supports growth and cues key aspects of *Pseudomonas aeruginosa* physiology. J Bacteriol 187:5267–77.

15. Son MS, Matthews WJ, Jr., Kang Y, Nguyen DT, Hoang TT. 2007. In vivo evidence of *Pseudomonas aeruginosa* nutrient acquisition and pathogenesis in the lungs of cystic fibrosis patients. Infect Immun 75:5313–24.

16. Hermansen GMM, Hansen ML, Khademi SMH, Jelsbak L. 2018. Intergenic evolution during host adaptation increases expression of the metallophore pseudopaline in *Pseudomonas aeruginosa*. Microbiology 164:1038–1047.

17. Imperi F, Tiburzi F, Visca P. 2009. Molecular basis of pyoverdine siderophore recycling in *Pseudomonas aeruginosa*. Proc Natl Acad Sci U S A 106:20440–5.

18. Yeterian E, Martin LW, Guillon L, Journet L, Lamont IL, Schalk IJ. 2010. Synthesis of the siderophore pyoverdine in *Pseudomonas aeruginosa* involves a periplasmic maturation. Amino Acids 38:1447–59.

19. Bleuel C, Grosse C, Taudte N, Scherer J, Wesenberg D, Krauss GJ, Nies DH, Grass G. 2005. TolC is involved in enterobactin efflux across the outer membrane of *Escherichia coli*. J Bacteriol 187:6701–7.

20. Andreini C, Banci L, Bertini I, Rosato A. 2006. Zinc through the three domains of life. J Proteome Res 5:3173–8.

21. Andreini C, Bertini I, Cavallaro G, Holliday GL, Thornton JM. 2008. Metal ions in biological catalysis: from enzyme databases to general principles. J Biol Inorg Chem 13:1205–18.

22. Poole K. 2005. Efflux-mediated antimicrobial resistance. J Antimicrob Chemother 56:20–51.

23. Furrer JL, Sanders DN, Hook-Barnard IG, McIntosh MA. 2002. Export of the siderophore enterobactin in *Escherichia coli*: involvement of a 43 kDa membrane exporter. Mol Microbiol 44:1225–34.

24. Crouch ML, Castor M, Karlinsey JE, Kalhorn T, Fang FC. 2008. Biosynthesis and IroC-dependent export of the siderophore salmochelin are essential for virulence of *Salmonella enterica* serovar Typhimurium. Mol Microbiol 67:971–83.

25. Tsutsumi K, Yonehara R, Ishizaka-Ikeda E, Miyazaki N, Maeda S, Iwasaki K, Nakagawa A, Yamashita E. 2019. Structures of the wild-type MexAB-OprM tripartite pump reveal its complex formation and drug efflux mechanism. Nat Commun 10:1520.

26. Li XZ, Plesiat P, Nikaido H. 2015. The challenge of efflux-mediated antibiotic resistance in Gram-negative bacteria. Clin Microbiol Rev 28:337–418.

27. Pearson JP, Van Delden C, Iglewski BH. 1999. Active efflux and diffusion are involved in transport of *Pseudomonas aeruginosa* cell-to-cell signals. J Bacteriol 181:1203–10.

28. Evans K, Passador L, Srikumar R, Tsang E, Nezezon J, Poole K. 1998. Influence of the MexAB-OprM multidrug efflux system on quorum sensing in *Pseudomonas aeruginosa*. J Bacteriol 180:5443–7.

29. Hirakata Y, Srikumar R, Poole K, Gotoh N, Suematsu T, Kohno S, Kamihira S, Hancock RE, Speert DP. 2002. Multidrug efflux systems play an important role in the invasiveness of *Pseudomonas aeruginosa*. J Exp Med 196:109–18.

30. Ramaswamy VK, Vargiu AV, Malloci G, Dreier J, Ruggerone P. 2018. Molecular Determinants of the Promiscuity of MexB and MexY Multidrug Transporters of *Pseudomonas aeruginosa*. Front Microbiol 9:1144.

31. Vega DE, Young KD. 2014. Accumulation of periplasmic enterobactin impairs the growth and morphology of Escherichia coli tolC mutants. Mol Microbiol 91:508–21.

32. Bleves S, Viarre V, Salacha R, Michel GP, Filloux A, Voulhoux R. 2010. Protein secretion systems in *Pseudomonas aeruginosa*: A wealth of pathogenic weapons. Int J Med Microbiol 300:534–43.

33. Hannauer M, Braud A, Hoegy F, Ronot P, Boos A, Schalk IJ. 2012. The PvdRT-OpmQ efflux pump controls the metal selectivity of the iron uptake pathway mediated by the siderophore pyoverdine in *Pseudomonas aeruginosa*. Environ Microbiol 14:1696–708.

34. Greenwald J, Hoegy F, Nader M, Journet L, Mislin GL, Graumann PL, Schalk IJ. 2007. Real time fluorescent resonance energy transfer visualization of ferric pyoverdine uptake in *Pseudomonas aeruginosa*. A role for ferrous iron. J Biol Chem 282:2987–95.

35. Jeong JY, Yim HS, Ryu JY, Lee HS, Lee JH, Seen DS, Kang SG. 2012. One-step sequence- and ligation-independent cloning as a rapid and versatile cloning method for functional genomics studies. Appl Environ Microbiol 78:5440–3.

36. Kaniga K, Delor I, Cornelis GR. 1991. A wide-host-range suicide vector for improving reverse genetics in gram-negative bacteria: inactivation of the blaA gene of *Yersinia enterocolitica*. Gene 109:137–41.

37. Richardot C, Plesiat P, Fournier D, Monlezun L, Broutin I, Llanes C. 2015. Carbapenem resistance in cystic fibrosis strains of *Pseudomonas aeruginosa* as a result of amino acid substitutions in porin OprD. Int J Antimicrob Agents 45:529–32.

38. Kessler E, Safrin M. 2014. Elastinolytic and proteolytic enzymes. Methods Mol Biol 1149:135–69.

39. Liberati NT, Urbach JM, Miyata S, Lee DG, Drenkard E, Wu G, Villanueva J, Wei T, Ausubel FM. 2006. An ordered, nonredundant library of *Pseudomonas aeruginosa* strain PA14 transposon insertion mutants. Proc Natl Acad Sci U S A 103:2833–8.

40. Tetard A, Zedet A, Girard C, Plesiat P, Llanes C. 2019. Cinnamaldehyde Induces Expression of Efflux Pumps and Multidrug Resistance in *Pseudomonas aeruginosa*. Antimicrob Agents Chemother 63.

41. Figurski DH, Helinski DR. 1979. Replication of an origin-containing derivative of plasmid RK2 dependent on a plasmid function provided in trans. Proc Natl Acad Sci U S A 76:1648–52.

