## Supplementary material for "Involvement of the *Pseudomonas aeruginosa* MexAB-OprM efflux pump in the secretion of the metallophore pseudopaline": Fig S1 & S2

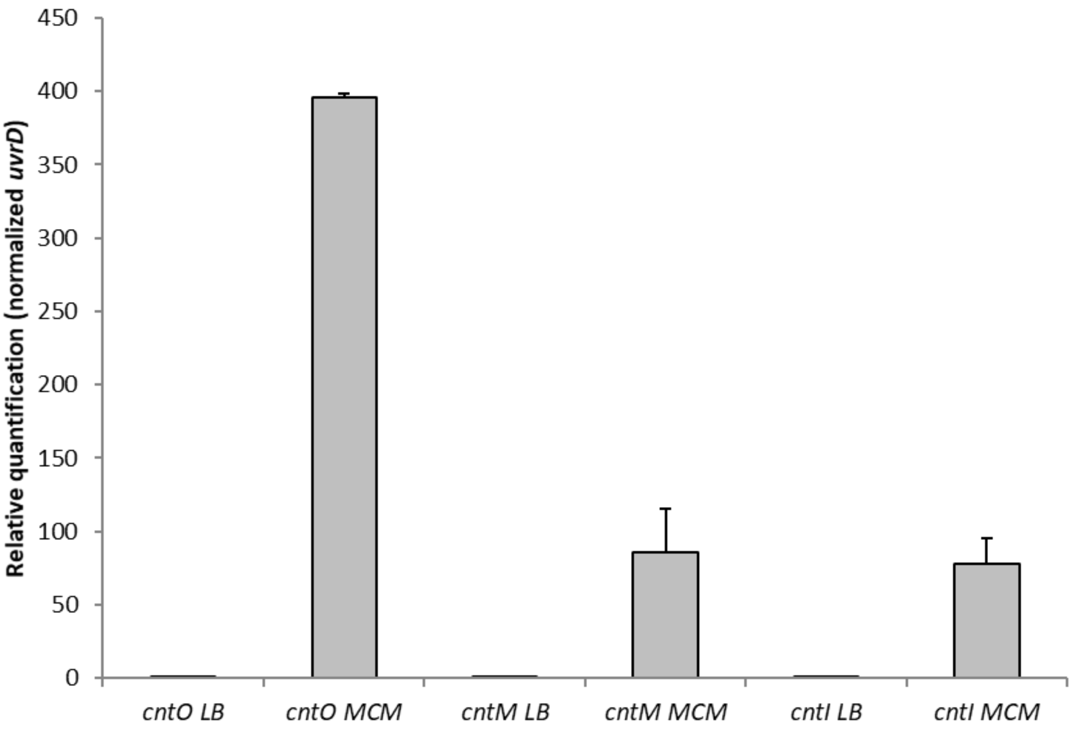


**Figure S1: Expression levels of *cntO, cntM* and *cntI* genes of the *cnt* operon in rich and minimal chelated medium.** Transcriptional activities were measured by RT-qPCR in PA14 wild type as described in the M&M section. The *uvrD* normalized transcriptional expressions of a technical duplicate of *cntO, cntM* and *cntI* in rich (LB) media and MCM in the wild type strain are presented.

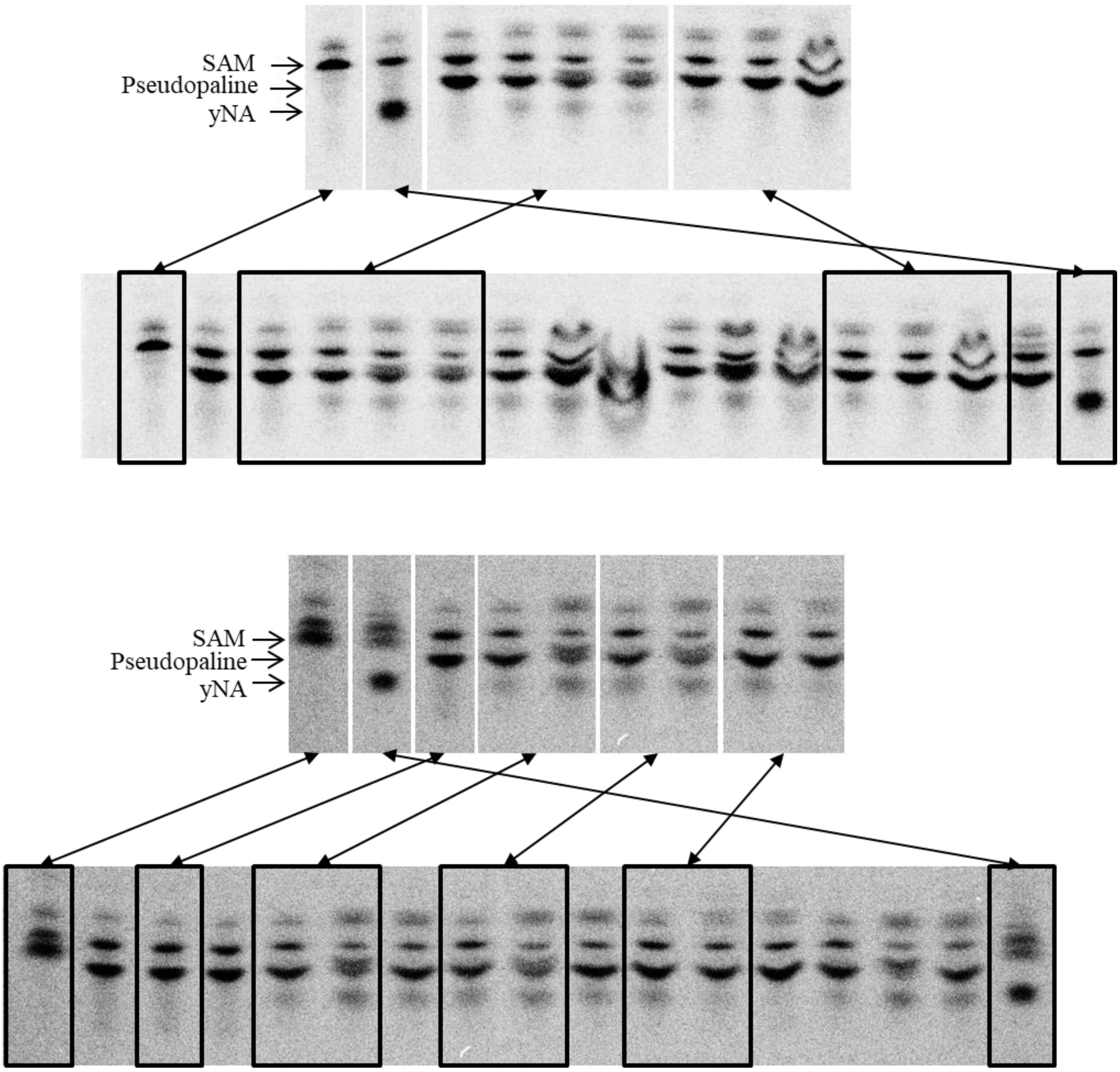


**Figure S2: original pictures of TLCs plates of *in vitro* pseudopaline degradation by PA14 cell lysates experiments presented figure 4.** Corresponding portions of TLCs used to assemble figure 4, extracted from unique TLC plates are indicated by black squares.
